# pyScoMotif: Discovery of similar 3D structural motifs across proteins

**DOI:** 10.1101/2023.08.27.554982

**Authors:** Gabriel Cia, Jean Marc Kwasigroch, Basile Stamatopoulos, Marianne Rooman, Fabrizio Pucci

## Abstract

**Motivation:** The fast and accurate detection of similar geometrical arrangements of protein residues, known as 3-dimensional (3D) structural motifs, is highly relevant for many applications such as binding region and catalytic site detection, drug discovery and structure conservation analyses. With the recent publication of new protein structure prediction methods, the number of available protein structures is exploding, which makes efficient and easy-to-use tools for identifying 3D structural motifs essential.

**Results:** We present an open-source Python package that enables the search for both exact and mutated motifs with position-specific residue substitutions. The tool is fast, accurate and suitable to run both on computer clusters and personal laptops. Successful application of pyScoMotif to catalytic site identification is showcased.

**Availability:** The pyScoMotif package can be installed from the PyPI repository and is also available at https://github.com/3BioCompBio/pyScoMotif. It is free to use for non-commercial purposes; for commercial purposes, please contact us.

## 1 Introduction

The function and biophysical properties of structured proteins are determined by the conformation adopted by their amino acid sequence, and in particular by the presence of specific local geometrical arrangements of subsets of residues. These motifs generally consist of few residues and typically correspond to binding regions or catalytic sites. They are known as 3D structural motifs and are generally conserved in homologous proteins across a wide range of organisms [1, 2]. A variety of *in-silico* tools to detect these motifs have been developed in the past decades, including brute-force methods [3, 4], geometric hash-based techniques [5, 6] and graph-based approaches [7, 8, 9, 10]. Recently, the Protein Data Bank (PDB) [11] made significant progress in this field by developing a novel and scalable structural motif search algorithm able to identify protein structures carrying a 3D motif that is similar to the queried motif [12]. The algorithm, which is inspired by the inverted index approach used in search engines, relies on the indexing of all the pairs of residues in a set of target protein structures. Once the index has been built, the search for similar motifs across the set of structures can be performed very fast by simply decomposing the query motif into a combination of residue pairs. Importantly, this approach does not require any pre-filtering or clustering of proteins at either the sequence or structure level, unlike some other methods [7, 3], thus enabling the discovery of 3D motifs even between evolutionary distant proteins.

The large-scale search for 3D structural motifs has recently attracted new interest thanks to the advances in protein structure prediction [13]. However, there is currently no tool available that is both efficient, flexible, and easy to use and to install. Here we present a Python package that implements a 3D structural motif search algorithm inspired by the PDB algorithm [12] with a series of improvements both in terms of search flexibility and user-friendliness.

## 2 Implementation

The pyScoMotif algorithm can be separated into two steps: indexing and motif search. We briefly describe the key elements of these two steps. Further details on the design and optimization choices as well as on the differences with respect to the PDB algorithm [12] are given in Supplementary Materials. A tutorial showing how to use pyScoMotif is available in our GitHub repository https://github.com/3BioCompBio/pyScoMotif.

### 2.1 Indexing

The objective of the indexing step is to extract and characterize the geometrical arrangement of all the residue pairs in a given set of protein structures, as shown in Figure 1. The resulting residue pair index is organized in such a way that searching the set of structures for all the occurrences of a residue pair with a specific geometry is very fast.

**Figure 1.**
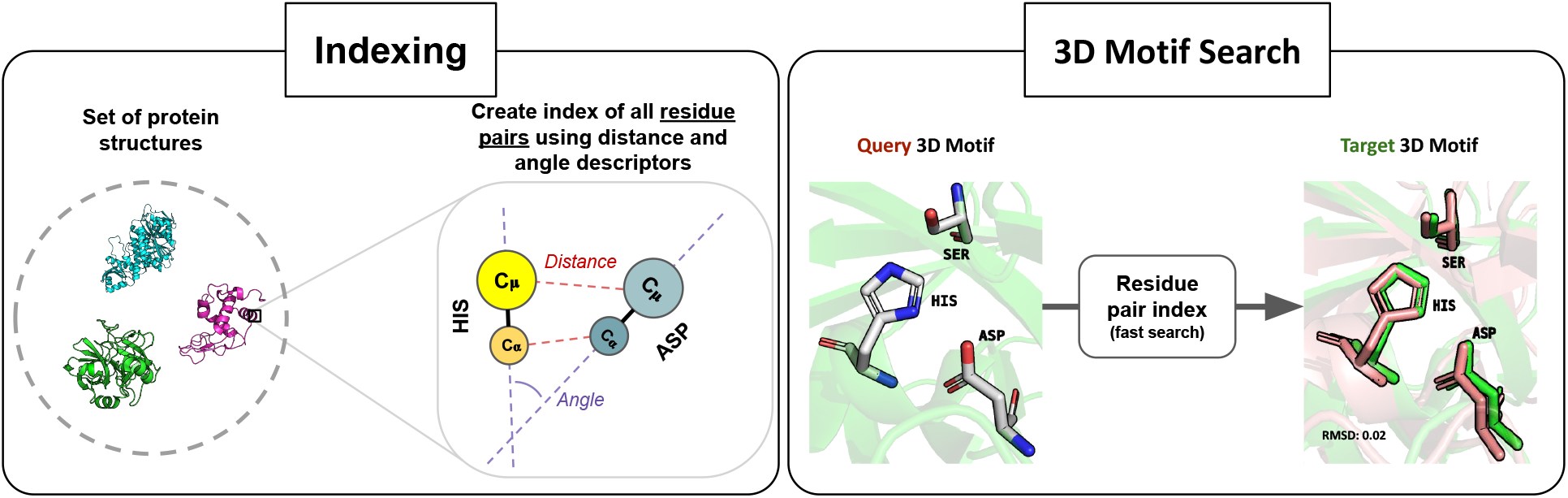
Schematic representation of the inverted index approach used in pyScoMotif to perform fast searches of similar 3D structural motifs in a set of protein structures. The first step involves pre-computing an index containing all the residue pairs in a given protein structure dataset based on their spatial arrangement (left). Using this pre-computed index, motif search can be performed very rapidly by decomposing the query motif into its residue pairs and loading them from the index (right). See section 2 for implementation details.

More precisely, given a set of protein structures, the indexing step generates a lookup table for each of the 210 possible residue pairs (*i*.*e*. Ala-Ala, Ala-Cys, …, Tyr-Tyr), with information about the geometrical arrangement of each occurrence of that residue pair. Three structural descriptors are used to yield a rotation-invariant geometrical description: the distance between the two *C*_*α*_ atoms; the distance between the two side chain centroids, noted *C*_*µ*_, calculated as the average of the coordinates of all the heavy side chain atoms; and the angle between the vectors 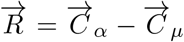 of the two residues. For the special case of Gly that has no heavy side chain atoms, we define *C*_*µ*_ = *C*_*α*_ and 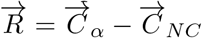, where *C*_*NC*_ is the midpoint between the amino N atom and the carboxyl C atom of Gly. Since structural motifs are by definition small sets of spatially close residues, we limit the indexation to residue pairs with a *C*_*α*_ *− C*_*α*_ distance *≤* 20 Å.

Each residue pair occurrence corresponds to a row in one of the lookup tables generated during indexation, and that row contains the identifier of the structure in which the pair occurs, the identifiers of the two residues, and the values of the three structural descriptors. Since the speed of the motif search is mainly limited by the loading speed of the lookup tables, it is advantageous to make them as small as possible [12]. Therefore, each residue pair lookup table is split into smaller tables that contain the data for a given range of values of the three structural descriptors. We used a 1Å bin size for the distance descriptors and a 20° bin size for the angle descriptor. These small lookup tables are implemented as simple CSV files. During the indexing process, each residue pair occurrence is appended at the end of its index file. Once all the occurrences of a residue pair have been processed, each index file is transformed into a pickled and compressed Pandas dataframe [14].

### 2.2 Motif search

The motif search step consists in finding all the 3D structural motifs in the set of indexed protein structures, called target motifs, that are similar to the motif given as input by the user, called query motif. Users must first provide a protein structure file (in PDB or mmCIF format) and the residue identifiers that make up the query motif. From this information, a fully connected, weighted, undirected graph is created to represent the motif, with nodes corresponding to residues and edges to the values of the three geometric descriptors of the residue pairs.

Searching the index tables for every single residue pair in the graph would be highly redundant and thus inefficient, especially for large graphs. Indeed, geometrical constraints between residue pairs have a transitive property (*i*.*e*. a constraint between residue pair (A,B) and (B,C) implies a constraint between residues A and C) which can be exploited to minimize the amount of data that is loaded from the index. Therefore, we prune graphs that have at least 4 nodes by applying Kruskal’s algorithm, which returns a minimum spanning tree (MST) of the graph that covers all the residues in the query motif. Then, for each residue pair in the MST, we load the index tables that match the values of the geometrical descriptors within the tolerance ranges specified by the user. As we process each residue pair, we iteratively check that the hits identified in each candidate structure are connected by an adge, otherwise the structure is discarded. Finally, the subset of structures that contain *all* the residue pairs in the MST is determined.

In the previous step, no check is performed regarding the *exact* connectivity of the residue pairs. As a result, some of the identified structures can have all the residue pairs in the correct geometrical arrangement but with different edge connections than in the query motif. To filter out these unwanted hits, we build the fully connected graph formed by all the identified residue pairs in each candidate structure and check for the presence of a subgraph that matches the query motif. The matching is performed using a subgraph monomorphism algorithm [15] (see Supplementary Material for details).

Finally, to calculate the root mean square deviation (RMSD) between the query motif and each target motif found, we superimpose the motifs using the fast quaternion-based superimposition algorithm [16] from Biopython [17]. Users can specify whether they want the RMSD to be calculated using the *C*_*α*_ coordinates, the *C*_*µ*_ coordinates, or both.

One of the main advantages of our implementation is that it allows users to specify different search strategies:

- Strict (by default): searches the index for target motifs that *exactly* match all the residues of the query motif.
- Position Specific Exchanges (PSE): searches the index for target motifs which may include mutated residues with respect to the query motif. Users have full control over which mutations should be allowed for each given residue in the query motif.
- Relaxed: similar to PSE search, but with possible mutations predefined based on physico-chemical amino acid properties: non-polar (G,A,V,L,I), polar (S,T,P,N,Q), sulfur-containing (M, C), positively charged (K,R,H), negatively charged (D,E) and aromatic (F,Y,W).
- Fully relaxed: extension of the relaxed search where residues can be mutated into any of the 19 other amino acids.

When performing searches with potential mutations, users can control the maximum number of simultaneous mutations, thus allowing them to search for a large number of mutated target motifs with a single search command.

## 3 Performance

We assessed pyScoMotif’s performance in terms of accuracy and speed and compared it with the PDB implementation [12], using the catalytic sites of serine protease, aminopeptidase and enolase, and the zinc-binding motif of the zinc-finger binding domain; details are given in Supplementary Material sections 4 and 5.

In terms of accuracy, the pyScoMotif and PDB motif search implementations return very similar results: when using each method’s default distance and angle thresholds and an RMSD threshold of 1 Å, we found an overlap of more than 95% between the results of the two methods on four different motifs (see Supplementary Materials section 5 for details). Note that we chose to use the side chain centroids *C*_*µ*_ rather than *C*_*β*_ atoms to represent residue sidechains, which makes our algorithm much more sensitive to correct side chain positioning than the PDB implementation.

pyScoMotif is highly parallelizable, thus making it highly efficient for both indexing and motif search. In terms of indexing speed, pyScoMotif is faster than the PDB implementation, as shown in Supplementary Material section 4. In terms of motif search speed, pyScoMotif is slightly slower than the PDB implementation, in large part due to the use of Python, which is inherently slower than Java. However, the bottleneck of the method is index creation, and pyScoMotif is able to perform motif searches in a few seconds, as shown in Table S2, which in absolute terms remains very fast.

## 4 Applications

We showcase a practical application of pyScoMotif: the identification of isoenzymes. Another application related to proteome-wide search for catalytic residues can be found in Supplementary Material section 6. We performed isoenzyme identification on alcohol dehydrogenase (ADH), an extensively studied class of enzymes that are well annotated in the Mechanism and Catalytic Site Atlas [18]. To check pyScoMotif’s sensitivity, we applied it to search for each ADH active site. We compared the results with the UniProt [19] annotations on all the structures in the index that belong to the ADH family.

In humans, alcohol metabolism is primarily achieved through the ADH enzyme family. There are seven ADH genes that code for seven ADH isozymes. The three genes ADH1A, ADH1B and ADH1C encode class I ADH isoenzymes, which have different substrate preferences and kinetics and are abundantly present in the liver [20]. As shown in Figure 2, the catalytic site is composed of three residues (Cys46, Cys174, His67) that bind the catalytic zinc ion, and two residues (Ser/Thr48, His51) that are involved in the catalytic reaction. In addition to the three class I ADH isoenzymes, humans encode four additional ADH classes [21]: class II (ADH4), class III (ADH5), class IV (ADH7) and class V (ADH6); although all the classes share a very similar catalytic site, some of them carry distinctive mutations.

**Figure 2.**
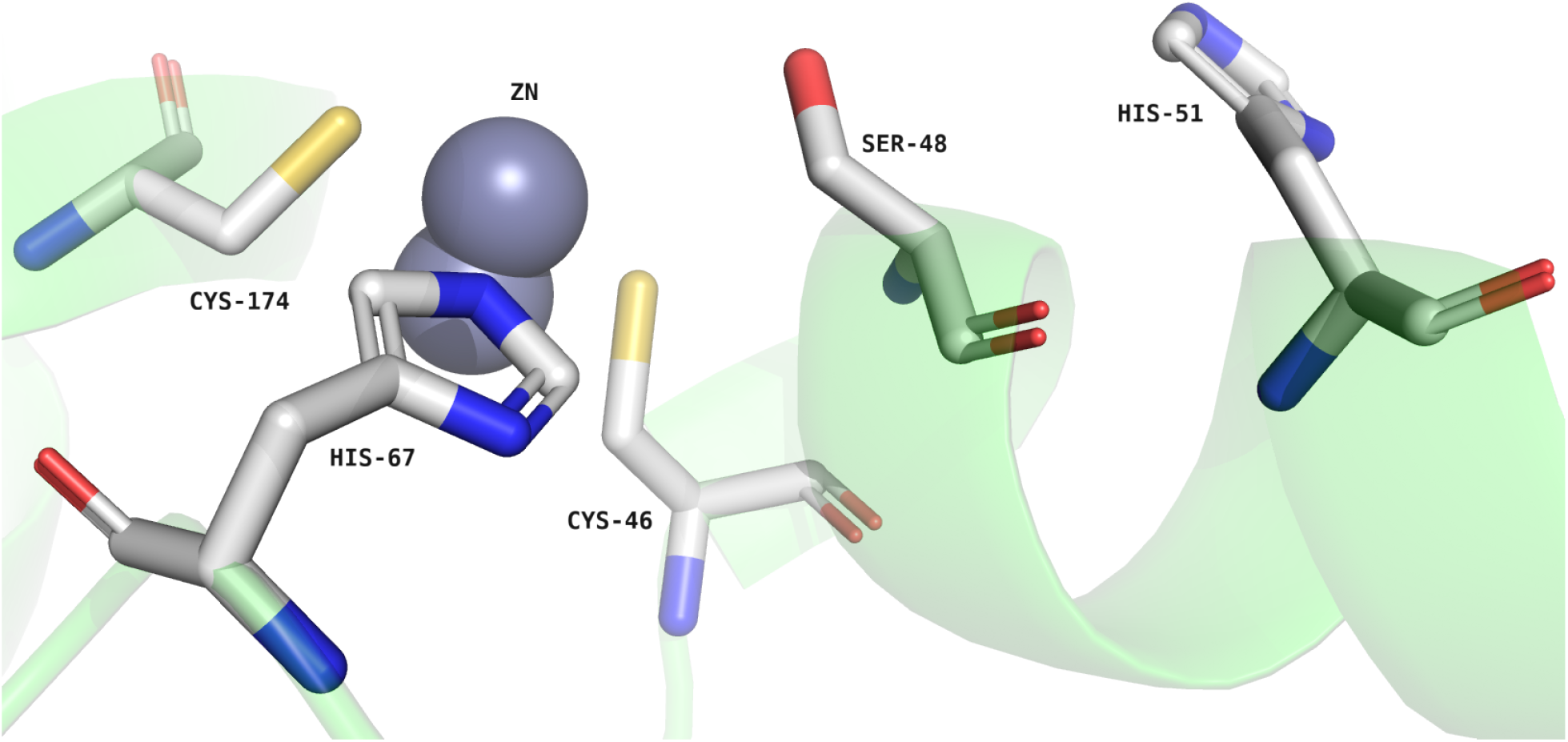
Catalytic site residues of the alcohol dehydrogenase structure PDB:1HSO. Generated with PyMOL [22].

We performed the search for motifs similar to the catalytic site of PDB:1HSO (the experimental protein structure of human ADH1A) on the human proteome, whose structures were taken from the AlphaFold database [13]. We used default search parameters, except for the residue type which was set to fully relaxed in order to detect the different ADH isoenzymes. pyScoMotif successfully reported the structures of the seven isoenzymes as the top hits, with RMSD values ranging from 0.21 for ADH1B (Uniprot ID: P00325) to 1.02 for ADH5 (Uniprot ID: P11766).

## 5 Discussion

This paper introduces pyScoMotif, an efficient, flexible and user-friendly Python implementation of a 3D structural motif search algorithm. The method allows searching for such motifs in large sets of protein structures in seconds, which otherwise could take hours using a brute force method. This is made possible by the use of an inverted index which organizes all the relevant structural information of the given set of protein structures in order to make 3D structural motif search very fast. The key elements of pyScoMotif are summarized below.

pyScoMotif is designed to be easy to install and use for researchers with no structural bioinformatics background. Its implementation balances speed and disk memory efficiency, thus allowing it to be used on both laptop computers and computer clusters. The code is highly parallelizable, which makes it suitable even for searches in large protein structure datasets. Since generating the index is pyScoMotif’s bottleneck in terms of time, we provide precomputed indexes of the full PDB as well as AlphaFold’s global health and human proteomes at http://babylone.ulb.ac.be/pyScoMotif/data/.

Another strength of pyScoMotif that sets it apart from other algorithms is its ability to perform flexible searches of the query motif. Indeed, motif search parameters such as tolerance thresholds of the geometrical arrangement of residues allow users to make their search as specific or loose as they wish. Moreover, users can search for possibly mutated versions of their motif by providing position-specific substitutions, and can even perform complex searches with multiple residues of the motif mutated simultaneously. The latter feature is particularly convenient as it allows users to search for a large number of similar motifs with a single command.

In summary, the rapid increase in 3D modelled protein 3D structures now enables large-scale structural analyses that would not have been possible previously. In this evolving context, pyScoMotif emerges as a valuable tool for researchers interested in analyzing newly available structural information and performing large-scale 3D structural motif searches.

