## Supplementary Material for "pyScoMotif: Discovery of similar 3D structural motifs across proteins"

###### **Index**

• 1. Introduction
• 2. Implementation details
• 3. pyScoMotif installation and usage
• 4. Performance
• 5. Benchmarking
• 6. Additional application example

### 1 Introduction

Three dimensional (3D) structural motifs are small ensembles of residues (typically less than 10) that have a structural or functional role, and tend to be well conserved across multiple species. The residues of these structural motifs are close in the 3D structure, but are generally distant in the protein sequence. This means that structure-based computational tools are needed to detect these motifs in protein structure databases. In addition, computational tools that can easily scale to databases containing hundreds of thousands of structures are needed, given the continuous increase in the number of available experimental and modeled protein structures [1, 2].

The Protein Data Bank (PDB) consortium [3] recently published a method [4] that is able to tackle these issues, which is available through the PDB website. Although the code is open source, it was not designed to be easily used as a standalone program, and it is thus difficult to use it to perform 3D structural motif searches on protein structures that are not included in the PDB. To fill this gap, we present pyScoMotif, a standalone Python implementation of an analogous algorithm with several improvements, which is very easy to use and can be installed with a simple pip install command.

#### 2 Implementation details

##### 2.1 Design and optimization

File input/output (IO) operations are the main limiting factor for both the indexing and motif search steps (which are described in the main manuscript), and one has to find a balance between speed and disk memory usage. Here we describe some of the important design and optimization trade-offs we made when implementing pyScoMotif.

The index lookup table files can consume a significant amount of disk space, especially when working with large protein structure databases. For example, indexing the entire PDB (accessed on 22/10/2022, $\sim 195,000$ PDB files) requires over 450GB of disk space. Our solution to this problem is simply to compress the index lookup table files, which reduces the disk memory footprint to 73GB while having a limited impact on the speed of the motif search.

Furthermore, the 210 residue pair combinations have to be processed in-dependently to effectively take advantage of file compression. This means that each structure file has to be read 210 times. Given that parsing PDB files was taking a significant amount of time and slowed down indexing, we decided to parse each structure file once and to store all relevant information in a Python pickle file, which is very fast to read. This approach made loading residue data $\sim 15$ times faster while having a minimal disk memory footprint. Note that these pickle files are also used during the motif search step to retrieve the coordinates needed to calculate the RMSD between each target motif and the query motif. Additionally, they make the index self-contained and portable since the original structure files used to create the index are not needed to perform motif searches.

Finally, note that each individual step of our algorithm (*i.e.* creation of the pickle structure files, indexing of the residue pairs, searching the index for residue pairs and RMSD calculation of each hit) can be parallelized, thus making our implementation scalable and efficient.

#### 2.2 Monomorphism and isomorphisms in motif search

To filter out the unwanted hits in the motif search step after the identification of all residues pairs in each candidate structure, we built their fully connected graph and checked for the presence of a subgraph that matches the user’s query motif.

The matching has to be performed using a subgraph monomorphism algorithm [5]. Subgraph isomorphism, which is the most commonly used graph morphism algorithm in chemoinformatics [6], would be incorrect if used in this instance. Indeed, graph edges do not correspond to chemical bonds, but simply to residue pair geometries, as shown on a hypothetical query and target motif graph in Figure S1. As a result, in certain cases, subgraph isomorphism would incorrectly reject hits that are in fact valid, whereas subgraph monomorphism correctly reports these hits.

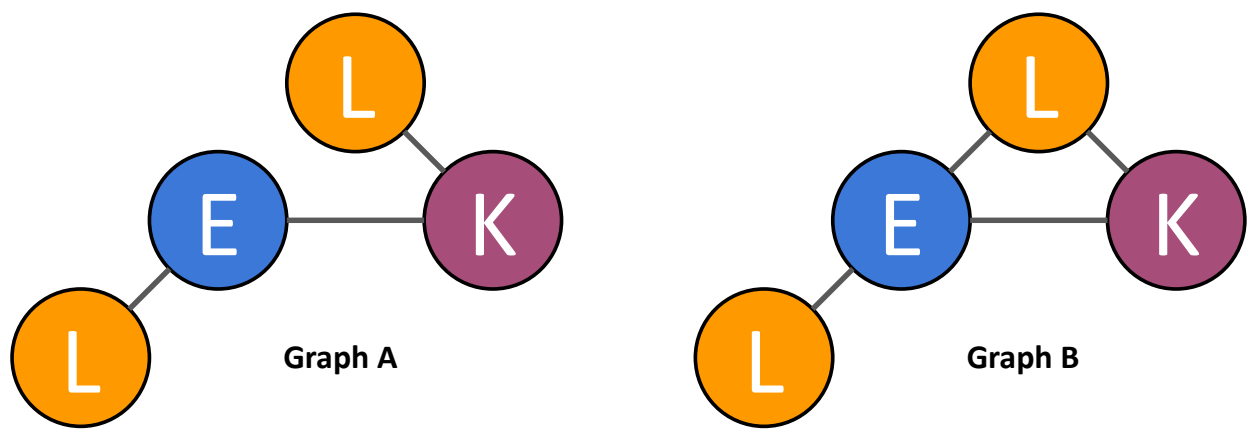

**Graphs A and B are monomorphic** (i.e. $A.nodes \equiv B.nodes$ AND $A.edges \subseteq B.edges$)
**but they are not isomorphic** (i.e. $A.nodes \equiv B.nodes$ AND $A.edges \equiv B.edges$)

Figure S1: Let A represent the pruned graph of a query motif, and B represent the graph of a candidate target motif. Subgraph isomorphism would incorrectly reject target motif B, while subgraph monomorphism would correctly return it as a hit. Indeed, edges do not correspond to chemical bonds, but simply to the occurrence of a residue pair in the correct geometrical arrangement, and in this example there are two occurrences of the residue pair EL that are in a correct geometry.

##### 3 pyScoMotif installation and usage

The pyScoMotif package can easily be installed through pip, the standard package manager for Python (`pip install pyscomotif`). It is implemented as a command line interface (CLI) with three commands. The full list of options and their detailed descriptions can be obtained by typing `pyscomotif <command-name> --help`.

1. The `create-index` command allows users to create the residue pair index that is required to perform a motif search. The command's signature is:

```
pyscomotif create-index --index_path=None --pattern=*.pdb
--compression=bz2 --n_cores=1 <database_path>
```

where `--index_path` is the full path of the directory that will contain the index files;

`--pattern` is the file extension pattern of the structure files to index;

`--compression` is the algorithm used to compress the index files;

`--n_cores` is the number of cores to use in parallel;

<database\_path> is the full path of the directory containing the structure files to index.

2. The `update-index` command allows users to update an existing index with new structures. The command's signature is:

```
pyscomotif update-index --pattern=*.pdb --n_cores=1
<database_path> <index_path>
```

where <index\_path> is the full path of the pyScoMotif index directory to update.

3. The `motif-search` command allows users to perform the search for 3D structural motifs in a given index of protein structures. The command's signature is:

```
pyscomotif motif-search --results_output_path=None
--residue_type_policy=strict --max_n_mutated_residues=1
--distance_delta_thr=2.0 --angle_delta_thr=30.0
--RMSD_atoms=CA+sidechain --RMSD_threshold=1.5 --n_cores=1
<index_path> <PDB_file> <motif>
```

where `--results_output_path` is the full path of the csv file where the pyScoMotif results will be saved;

`--residue_type_policy` is an option to control whether mutated versions of the query motif should also be searched, which can be set to either strict (default), relaxed, fully\_relaxed or a position-specific substitutions, in JSON format (see end of the "Motif search" subsection for details about each option);

`--max_n_mutated_residues` is the maximum number of simultaneous mutations that are allowed when generating mutated motifs through the `--residue_type_policy` option;

`--distance_delta_thr` is the allowed distance tolerance (in Å) for considering two pairs of residues as similar;

`--angle_delta_thr` is the allowed angle tolerance (in degrees) for considering two pairs of residues as similar;

`--RMSD_atoms` allows users to decide whether the coordinates of $C_\alpha$ atoms, $C_\mu$ atoms or both are used to calculate the RMSD;

`--RMSD_threshold` allows users to set the maximum RMSD value of the similar motifs reported in the output csv table;

<PDB\_file> is the full path of the structure file containing the query motif;

<motif> is a list of space-separated residue identifiers contained in the query motif.

For more information about these commands and a detailed tutorial on how to use our tool, see the pyScoMotif guide at https://github.com/3BioCompBio/pyScoMotif. Note that pre-built indexes of the full Protein Data Bank, the AlphaFold2 global health proteomes and the human proteome are available at http://babylone.ulb.ac.be/pyScoMotif/data/.

#### 4 Performance

This section shows the speed and disk memory consumption for the index creation and motif search commands. All the values were obtained on a computer with a 3.8GHz AMD Threadripper CPU, a 512GB NVMe SSD with read/write speed of 3,500/2,300 Mb/s, and 32GB of 3200MHz DDR4 RAM, using Python 3.10.6. The timings were obtained using the Unix `$time` command, and the pyScoMotif commands were all parallelized on 12 cores.

| Database | Database size (GB) | Index disk memory (GB) | Index creation speed |
| --- | --- | --- | --- |
| *S.Cerevisiae* AF proteome | 0.5 | 1.7 | 13 min 30 sec |
| *H.Sapiens* AF proteome | 2.1 | 6.1 | 1 hour 7 min |
| *G.Max* AF proteome | 3.2 | 8.8 | 1 hour 33 min |
| Full PDB | 41.4 | 73 | 20 hours 30 min |

Table S1: Disk memory consumption and execution time of pyScoMotif to index the proteomes of *S. Cerevisiae*, *H. Sapiens* and *G. Max*, which were downloaded from the AlphaFold (AF) database [1], as well as of the full PDB (22/10/2022, $\sim 195,000$ PDB files). Database sizes corresponds to gzip compressed PDB files.

Table S1 reports the index creation speed and disk memory usage on three proteomes of which the protein structures were available in the AlphaFold database [1], *i.e.* *S. Cerevisiae*, *H. Sapiens* and *G. Max*, as well as on the full PDB; note that these timings are dependent on the hardware used. The results show that pyScoMotif’s speed scales linearly with the size of the database, as expected. It only took 20 hours 30 min to index the entire PDB ($\sim 195.000$ PDB files), whereas the PDB implementation paper reports that it took 3 days and 11h to index an earlier version of the PDB dataset ($\sim 170.000$ PDB files) using the same number of parallel cores, which shows pyScoMotif’s superior indexing speed.

We also evaluated pyScoMotif’s motif search speed on the set of protein catalytic sites that were evaluated in the paper presenting the PDB algorithm ([4], Table 1), which include: serine protease (PDB:4CHA - His:B57, Asp:B102, Ser:C195), aminopeptidase (PDB:1LAP - Lys:A250, Asp:A255, Asp:A273, Asp:A332, Glu:A334), zinc finger domain (PDB:1G2F - Cys:F207, His:F225, His:F229) and enolase (PDB:2MNR - Lys:A164, Asp:A195, Glu:A221, Glu:A247, His:A297). We evaluated pyScoMotif’s performance both on human proteome structures taken from the AlphaFold database and the full PDB. Also, in order for values to be comparable with those of the PDB algorithm paper, we used their distance ($1\text{\AA}$), angle ($20^\circ$) and RMSD ($1\text{\AA}$) thresholds on all the following analyses.

| Motif searched | AF *H.Sapiens* |  | Full PDB |  |
| --- | --- | --- | --- | --- |
|  | Execution speed | Hits | Execution speed | Hits |
| Serine protease | 1.8 sec | 112 | 4.8 sec | 3,027 |
| Aminopeptidase | 1.8 sec | 2 | 3.9 sec | 390 |
| Zinc finger domain | 3.4 sec | 6,207 | 2.9 sec | 1,098 |
| Enolase | 1.8 sec | 1 | 3.4 sec | 284 |

Table S2: Execution speed and hits reported by pyScoMotif on four 3D motifs against the AlphaFold (AF), *H.Sapiens* proteome and the full PDB.

The results in Table S2 show that motif search takes few seconds for any motif, which is much faster than any brute force approach. In addition, search speed scales sub-linearly with database size, making it particularly efficient on very large datasets. Indeed, despite the full PDB database being 20 times larger than the AF human proteome database, the search speed of the serine protease, aminopeptidase and enolase catalytic sites only increased by a factor of about 2, as shown in Table S2.

The comparison of pyScoMotif’s search speed on the full PDB with the values reported in the PDB implementation paper ([4], Table 1) indicates that pyScoMotif is about 5 times slower. Note that this is to be expected given pyScoMotif is implemented in Python while the PDB implementation is in Java, which is known to be faster.

#### 5 Benchmarking

We performed two benchmark experiments to analyse and compare the hits detected by pyScoMotif and the PDB implementation:

1. The first experiment evaluates the overlap between the hits predicted by both methods on the four motifs from Table S2, which were searched against the full PDB.
2. For the second experiment, we implemented an exhaustive full atom motif search algorithm. We calculated the Positive Predictive Value (PPV) and False Negative Rate (FNR) of pyScoMotif and the PDB implementation on the same four motifs discussed above.

##### 5.1 Overlap between pyScoMotif and the PDB results

We benchmarked pyScoMotif against the PDB implementation [4] by comparing the hits returned by both methods on the four motifs listed in Table S2. The PDB hits were obtained by running each motif through the PDB online structure motif search on the same set of indexed structures (*i.e.* experimental PDB structures released before 22/10/2022).

To evaluate the similarity of the results of both methods, we calculated the number of hits identified by both methods over the number of hits that are found by at least one of the methods (intersection over union). When running pyScoMotif with the same distance, angle and RMSD thresholds than the PDB implementation, we found an intersection over union of 82%, 98%, 65% and 83% for the serine protease, aminopeptidase, zinc finger domain and enolase motifs, respectively. The observed differences are simply a consequence of the different structural descriptors used by the two methods to capture side chain information. Indeed, pyScoMotif uses the $C_\mu$ pseudoatoms (centroids of heavy side chain atoms), whereas the PDB implementation uses the $C_\beta$ atoms. Therefore, pyScoMotif is much more sensitive to the correct positioning of all the residues’ side chains than the PDB implementation. This is a thoughtful choice as we believe the side chain positioning is crucial in 3D structure motifs.

As shown in Figure S2, pyScoMotif is able to detect target motifs where the side chain orientations are conserved (Figure S2a), but does not report target motifs where at least one of the side chains is in a significantly different orientation (Figure S2b, see ASP-102). Conversely, the PDB implementation is very sensitive to the conservation of the $C_\beta$ position, and thus target motifs where this atom is significantly shifted but where the side chain is generally conserved are not reported (Figure S2a), whereas it is able to report target motifs where the $C_\beta$ position is conserved but the side chain is not (Figure S2b).

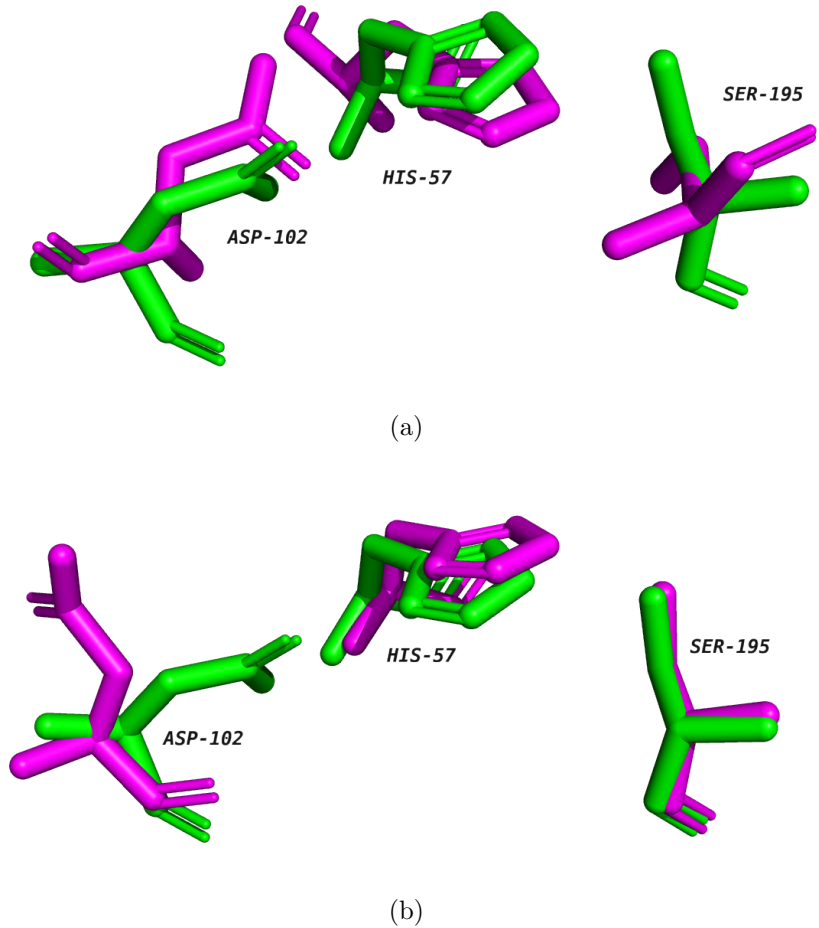

Figure S2: Examples of missed target motifs by each method on the serine protease catalytic site. The true catalytic site residues are in green, while the missed target motifs are in magenta. **(a)** Target motif missed by the PDB implementation but detected by pyScoMotif. **(b)** Target motif missed by pyScoMotif but detected by the PDB implementation. Pictures generated with PyMOL [7].

We also verified that enlarging the pyScoMotif distance and angle thresholds to their default values (*i.e.* 2.0 Å and 30° respectively) but keeping the 1 Å RMSD threshold, results in an overlap between the pyScoMotif and PDB implementation hits of $\geq 95\%$ on the four motifs, and even reached 100%

of the hits for aminopeptidase. In summary, both methods return similar results for a given RMSD threshold.

#### 5.2 Benchmarking against an exhaustive full atom motif search algorithm

Both pyScoMotif and the PDB implementation index use a compressed representations of amino acids, which are encoded using only two coordinates: the ($C_\alpha$), which represents the main chain, and the $C_\beta$ or $C_\mu$, which represent the side chain. This could result in some incorrect hits or misses with respect to an exhaustive atom based motif search algorithm.

To evaluate this, we implemented such an exhaustive and time-consuming motif search algorithm that searches 3D structural motifs using subgraph monomorphism on full atom graphs, as opposed to the simplified graphs used by both pyScoMotif and the PDB implementation. We used the four motifs from Table S2 used above which we searched against the AF human proteome. Choosing the exhaustive full atom search algorithm results as the ground truth, we calculated the PPV and FNR of both methods, which are reported in Table S3.

Overall, the results show that pyScoMotif and the PDB implementation have very similar PPV and FNR scores across all four motifs, with the exception of the FNR on the zinc finger domain motif where the PDB implementation performs slightly better than pyScoMotif (0.10 vs 0.17, respectively). This can be explained by the fact that both the PDB implementation and our full atom motif search algorithm calculate the RMSD between query and target motifs using full atom representations, unlike pyScoMotif which calculates its RMSD values using only $C_\alpha$ and $C_\mu$ coordinates. The PDB implementation has thus a slight advantage in this benchmark, but this comes at the cost of storing a large database with the atomic information of all indexed structures. Note that running the PDB implementation with RMSD values computed using the $C_\alpha$ and $C_\beta$ coordinates leads to lower PPV values (see Table S3), especially for the serine protease motif where it obtains a PPV of 0.54, which is significantly worse than pyScoMotif.

| Motif searched | pyScoMotif ($C_\alpha$ & $C_\mu$ RMSD) | | PDB (full atom RMSD) | | PDB ($C_\alpha$ & $C_\beta$ RMSD) | |
| --- | --- | --- | --- | --- | --- | --- |
|  | PPV | FNR | PPV | FNR | PPV | FNR |
| Serine protease | 0.97 | 0.04 | 1.00 | 0.03 | 0.54 | 0.03 |
| Aminopeptidase | 1.00 | 0.00 | 1.00 | 0.00 | 1.00 | 0.00 |
| Zinc finger domain | 0.88 | 0.17 | 0.90 | 0.10 | 0.86 | 0.10 |
| Enolase | 1.00 | 0.00 | 1.00 | 0.00 | 1.00 | 0.00 |

Table S3: Positive Predictive Value (PPV = TP / (TP+FP)) and False Negative Rate (FNR = FN/(FN+TP)) of pyScoMotif and the PDB implementation when taking an exhaustive full atom motif search algorithm as the ground truth.

In summary, it is noteworthy that pyScoMotif obtains very similar or only marginally lower performances with its compressed and memory-efficient representation of amino acids to compute the RMSD. Moreover, it is important to keep in mind that the concept of absolute or ground truth in the context of motif search is ill defined and problem dependant. For example, searching for both active and inactive conformations of a catalytic site or just for one of the two conformations results in different ground truths. Note, in addition, that inaccuracies in the exact positioning of all side chain atoms, a known issue in structure modelling, can be at least partly alleviated by using pyScoMotif’s average side chain centroids.

#### 6 Additional application example

Here we show an application of pyScoMotif for the detection of catalytic sites across proteomes. We used the catalytic sites of tryptophan synthase as an example. Bacteria, fungi and plants are able to biosynthesize tryptophan through the tryptophan synthase enzyme. Animals on the other hand do not express tryptophan synthase and instead rely on diet to obtain this essential amino acid. The tryptophan synthase biological unit is a $2\alpha$-$2\beta$ tetramer structure [8], and each $\alpha$ and $\beta$ subunit has a catalytic site that performs a specific catalytic reaction.

As shown in Figure S3, the $\alpha$ chain catalytic site has seven residues (Gly49, Asp60, Tyr102, Tyr175, Thr183, Ser233, Ser235), and the $\beta$ chain catalytic site has six residues (Lys87, Thr110, Thr190, Ser235, Asn236 Ser377). As a positive test, we checked for the presence of the two catalytic sites in *H. Pylori* (bacteria), *S. Cerevisiae* (fungi) and *G. Max* (plant), and used the human proteome as a negative check. The proteome of each respective organism was downloaded from the AlphaFold database [1] and used to build a pyScoMotif index. We performed the searches of similar motifs using the $\alpha$ and $\beta$ chain catalytic sites of PDB:1A50, the tryptophan synthase structure of *S. typhimurium*.

• For *H. Pylori*, there is one $\alpha$ and one $\beta$ chain structure in the AlphaFold proteome, and pyScoMotif correctly reported these two hits only.
• For *S. Cerevisiae*, there is only one heterodimeric structure in the AlphaFold proteome, which carries a Ser190Ile mutation on the $\beta$ chain, and pyScoMotif correctly reported that structure when searching for the catalytic sites with the right position-specific exchange.
• For *G. Max*, there are two $\alpha$ and four $\beta$ chain structures in the AlphaFold proteome. For the $\alpha$ chain catalytic site, both structures feature a Ser233Val mutation as well as a moderate shifting of Glu49 side chain position. pyScoMotif correctly reported both of them when using the appropriate position-specific exchange and raising the angle tolerance to $45^\circ$ to account for the shifting of Glu49. For the $\beta$ structures, mutations Thr190Ser and Ser377Cys are present in two of the structures as well as a moderate shifting of the Lys87 side chain position in one structure, and pyScoMotif correctly reported the four structures when using the appropriate position-specific exchange, setting the maximum number of mutated residues to two and raising the angle tolerance to $45^\circ$ to account for the shifting of Lys87.
• For the human proteome, there was no tryptophan synthase structure, and pyScoMotif correctly reported zero hits for both catalytic sites.

This example shows another key strength of pyScoMotif, *i.e.* how the use of the motif search options such as position specific exchanges make the search for mutated motifs very easy.

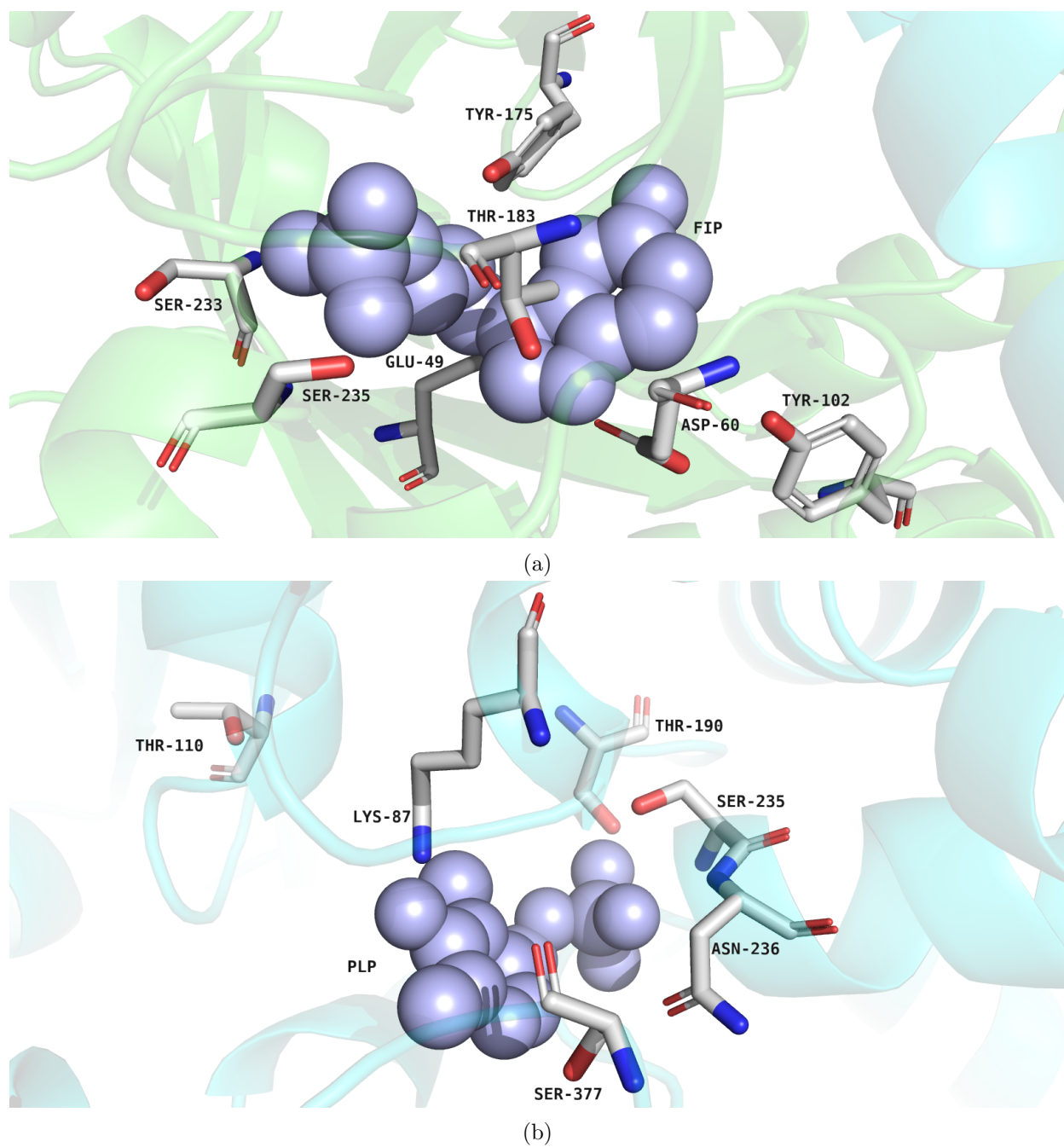

Figure S3: Catalytic site residues of the $\alpha$ (top) and $\beta$ (bottom) subunits in the tryptophan synthase structure PDB:1A50. Pictures generated with PyMOL [7].

#### References

- [1] Mihaly Varadi, Stephen Anyango, Mandar Deshpande, Sreenath Nair, Cindy Natassia, Galabina Yordanova, David Yuan, Oana Stroe, Gemma Wood, Agata Laydon, et al. AlphaFold Protein Structure Database: mas-

- sively expanding the structural coverage of protein-sequence space with high-accuracy models. *Nucleic acids research*, 50(D1):D439–D444, 2022.
- [2] Zeming Lin, Halil Akin, Roshan Rao, Brian Hie, Zhongkai Zhu, Wenting Lu, Nikita Smetanin, Robert Verkuil, Ori Kabeli, Yaniv Shmueli, Allan dos Santos Costa, Maryam Fazel-Zarandi, Tom Sercu, Salvatore Candido, and Alexander Rives. Evolutionary-scale prediction of atomic level protein structure with a language model. *bioRxiv*, 2022.
- [3] Helen M Berman, John Westbrook, Zukang Feng, Gary Gilliland, Tala-pady N Bhat, Helge Weissig, Ilya N Shindyalov, and Philip E Bourne. The protein data bank. *Nucleic acids research*, 28(1):235–242, 2000.
- [4] Sebastian Bittrich, Stephen K Burley, and Alexander S Rose. Real-time structural motif searching in proteins using an inverted index strategy. *PLoS computational biology*, 16(12):e1008502, 2020.
- [5] Aric Hagberg, Pieter Swart, and Daniel S Chult. Exploring network structure, dynamics, and function using NetworkX. Technical report, Los Alamos National Lab.(LANL), Los Alamos, NM (United States), 2008.
- [6] John W Raymond and Peter Willett. Maximum common subgraph isomorphism algorithms for the matching of chemical structures. *Journal of computer-aided molecular design*, 16:521–533, 2002.
- [7] Warren L DeLano et al. Pymol: An open-source molecular graphics tool. *CCP4 Newsletter on protein crystallography*, 40(1):82–92, 2002.
- [8] CC Hyde, SA Ahmed, EA Padlan, E Wilson Miles, and DR Davies. Three-dimensional structure of the tryptophan synthase alpha 2 beta 2 multienzyme complex from salmonella typhimurium. *Journal of Biological Chemistry*, 263(33):17857–17871, 1988.
